# A Markov chain model of cancer treatment

**DOI:** 10.1101/2021.06.16.448669

**Authors:** Péter Bayer, Joel S. Brown, Johan Dubbeldam, Mark Broom

**Author notes:** We thank Nathaniel Mon Père for sharing his ideas with us and for assisting in conducting simulations.

## Abstract

This paper develops and analyzes a Markov chain model for the treatment of cancer. Cancer therapy is modeled as the patient’s Markov Decision Problem, with the objective of maximizing the patient’s discounted expected quality of life years. Patients choose the number of treatment rounds they wish to administer based on the progression of the disease as well as their own preferences. We obtain a powerful analytic decision tool by which patients may select their preferred treatment strategy. In a second model patients may make choices on the timing of treatment rounds as well. By delaying a round of therapy the patient forgoes the gains of therapy for a time in order to delay its side effects. We obtain an analytic tool that allows numerical approximations of the optimal times of delay.

## 1 Introduction

Cancer treatment faces many unique challenges. Arguably the most important one is that the available therapies against the metastatic disease produce very high failure rates. As such, since outright cure is unlikely, and the therapies themselves are invasive, costly, and often come with a significant reduction to the patient’s quality of life, metastatic cancer treatment comes with difficult dilemmas that require tradeoffs between curing the patient in terms of maximizing the probability of success against caring for the patient in terms of their well-being. Preserving a high quality of life, maximizing the probability of recovery, or the patient’s life expectancy cannot always be achieved through the same treatment strategy. Resolving these dilemmas in practice is further constrained by the necessarily high legal standards of medicinal practice and the treatments’ economic and budgetary considerations, as well as patient autonomy.

In this paper we provide a theoretical foundation to formally capture these dilemmas. By employing mathematical tools, particularly dynamic optimization, statistics and game theory, we build a model of cancer treatment by which these dilemmas can be explicitly addressed. We wish to make no pretense that our model captures all possible such dilemmas, or that its predictions represent the uniquely correct way of resolving the ones that we do consider; instead our intention is to introduce methods and concepts by which the discussion surrounding them can be advanced.

Survival time remains the prevailing measure of success in cancer therapy. Due to the unambiguity and availability of data it is the least controversial and most accessible metric. Mathematical models of cancer therapy often report on their proposed regimens’ effects on (simulated) survival or progression time. Clinical trials of new drugs and methods of delivery are similarly evaluated on this basis. Yet, there is reason to believe that oncologists and patients do not make treatment decisions to maximize survival time. In particular, decisions to refuse therapy are often influenced by concerns over quality of life (Shumay et al., 2001) and cure probability (Frenkel, 2013) possibly at the expense of expected survival time. While the prevailing response to such decisions had been a call for oncologists to “better communicate” with their patients, whether the prescribed therapy indeed aligns with the patient’s objectives is not so clear. In particular, patients who refuse therapy at times report no worse quality of life than those who complete it (Gilbar, 1991).

Even if a positive definition could be given that defines the goals and aims of cancer therapy with respect to improving the patient’s health outcomes, patient autonomy means that treatment decisions also take into account the patient’s own wishes. What is at hand, therefore, is a strategic choice of treatment strategy that is made in regards to a combination of objective concerns relating to disease prognosis and subjective ones relating to the patient’s personal preferences. Moreover, as cancer therapy is a long process with choices having to be made and re-made in response to the progression of the disease as well as the consequences of past choices, models that seek to inform cancer therapy need to be dynamic and allow for multiple points of decision making.

The tools and concepts of game theory and decision theory have proven extremely valuable in cancer research. The objective has been to utilize game theory’s insights in understanding the eco-evolutionary dynamics of cancer. The practical application of this research direction is thus, first, to calibrate the parameters (doses, timing, duration) of existing therapy regimens (see e.g. adaptive therapy, Gatenby et al., 2009) and, second, to find new points of attack against the disease in search of new therapy regiments.

One development towards the former branch is the concept of viewing cancer therapy as a game played between the disease and the treating physician (Orlando et al., 2012). A useful framework is to model the game as a leader-follower (Stackelberg-)game with the treating physician as a strategic decision maker and cancer as a reactive and adaptive player, its strategies being a consequence of it undergoing evolution by natural selection to the environment influenced by the physician’s chosen treatment strategies (Staňková et al., 2019). The key insight of this analogy is to identify the benefits that the physician can realize by assuming the leader role in the game and use the information about cancer’s the possible reactions to their advantage. Instead, we often observe physicians in the reactive role and following a prescribed or standard treatment strategy, changing only after observing a new strategy from the disease.

We advance this thread of the literature by viewing the game as a Markovian process. Such processes have an established application in cancer (Kay, 1986; Andersen et al., 1991). In Markovian models of cancer, all relevant information regarding the prognosis of the patient is encoded in health states, usually including a healthy state, various states of disease progression, and a death state. The patient transitions between these states according to a stochastic process. The transition probabilities of such models may be calibrated from cohort data (Duffy et al., 1995) for simulations of likely disease progression. The resultant toolkit has applications in both medicine (Llorca et al., 2001) and health economics (Le Lay et al., 2007).

Crucially, in Markovian models, the transition probabilities are assumed to depend only on the current state of the patient, not on previous disease history. This is both a simplifying and a limiting assumption that presents a modeling challenge: too few health states may obscure progression-relevant patient information while having too many health states is impractical for applications and may fail to produce insight that can be generally applied to a large cohort of patients. To resolve this, Cooper et al. (2003, 2004) introduced a small number of payoff-states (responsive, stable, progressive, dead), but allowed for changing transition dynamics between them based on the length of the treatment, measured in the number of treatment cycles.

To this existing framework we add the element of choice by the patient.^1^ Markov decision processes (MDPs) (Bellman, 1957) combine the tools of stochastic processes and decision theory. In this model the Markovian transition probabilities depend upon both the current state and the strategy of a payoff-maximizing decision maker. The patient receives payoffs, measured in quality adjusted life years (QALYs), from spending time in states, with more healthy states giving higher payoffs. The tension in these problems is introduced when the decision-maker faces a choice between strategies that lead to immediate payoff gains and strategies that lead to better future prospects but at the cost of foregoing immediate gains. These trade-offs are also highly relevant in the choice of cancer therapy; the choice of taking therapy involves an investment by the patient, both in financial and in QALY terms, in the hopes of a higher probability of cure and greater life expectancy. Under the classic results of MDP literature (Blackwell, 1962, 1965), if the decision-maker’s objectives can be represented by discounting future expected payoffs and the set of states is finite, then an optimal policy will exist and is generically unique (Ortega-Gutiérrez et al., 2016).

In this paper we use MDPs to model the novel idea of the game between the physician and the disease in a Markovian environment. By this approach the game is reduced to a problem with a single strategic decision maker, the patient. We treat the evolutionary processes of cancer as an exogenous and stochastic element, whose behavior, conditional on the selected treatment strategy, can be estimated from cohort data. We introduce exponential discounting to model a preference for earlier QALYs over later ones. As a treatment strategy will always exist that maximizes discounted expected QALYs, we are able to derive optimal treatment strategies.

We first place the focus on the duration of treatment. The patient’s payoff is the difference between their QALYs and the cost of the treatment. The main tension in our model is the trade-off between continuing with the treatment and bearing the cost in hopes of a higher cure probability and/or longer life expectancy, or abandoning treatment. A complicating factor is the adaptive dynamics of cancer. As the patient progresses through rounds of treatment, cancer’s responsiveness to the therapeutic agent changes. Following Cooper et al. (2003) our model has an infinite series of health states (other than the absorbing ‘cured’ and ‘death’ states) with the *i*th clone of a health state representing the patient after *i* rounds of medication. Each clone of the same health state offers the same QALYs as the original but may have different transition probabilities to other states. After another round of therapy, unless the patient moves to either of the two absorbing states, he or she moves up to the *i* + 1th clone, thus the next round of therapy will happen under different transition rates. This model allows us to derive conditions on the number of rounds a payoff-maximizing patient takes.

From this model we are able to derive efficient methods to evaluate treatment strategies of different duration. Under two monotonicity conditions on the parameters, the patient’s best treatment strategies may be derived analytically: in this case a myopic treatment plan, i.e. administer therapy if and only if one more round is better than no more rounds, identifies the globally optimal treatment strategy. In particular, if the patient’s likelihood of recovery is not increasing with each new dose of treatment, an assumption that is motivated both by the onset of resistance to therapy as well as observed outcomes of cancer therapy, there will exist a unique payoff-maximizing duration of therapy, beyond which patients lose expected QALYs due to overtreatment. We simulate this effect and show that, while the ex-ante expected payoff loss of overtreatment may be marginal due to time-discounting and the cohort’s attrition up to the time when overtreatment is reached, the realized payoff loss for patients who do reach that stage is substantial.

In a second model we internalize effects of treatment to the patient’s QALYs. As cytotoxic therapy of cancer is often highly toxic for the patient, a major constraint in the timing of doses, and, as discussed, one of the main incentives to refuse or abandon therapy, is the lost quality of life under therapy. We thus make this element explicit in our model; the payoff of the patient depends upon their current health state and the current level of toxicity. We assume that each round of therapy adds to the patient’s toxicity level which depreciates over time. When taking therapy the QALY-cost of therapy is not instantaneous, as in our base model, but is incurred continuously. This changes the game compared to our base model in two ways. First, the cost of therapy becomes conditional on its outcome; surviving patients have to bear the QALY reduction longer, while patients who are not cured may have to resort to taking on additional QALY reductions. Second, patients are afforded the option to reduce the QALY-cost of therapy by postponing it, allowing their level of toxicity to depreciate before taking on additional QALY reductions. However, by doing so they also postpone any benefits of therapy to their recovery, introducing another source of tension to the model.

Under classical MDPs, in which the decision maker’s payoffs depend only on their current state, in optimum, the decision maker’s choice in any given state does not change until he or she transitions to the next state. This, however, is not a sensible conclusion for cancer therapy as the few health states usually fail to capture all relevant patient information, thus the optimal course of therapy may change before the patient transitions to a new health state. Our model of toxicity accounts for this as well, as the patient’s choice of therapy is allowed to be dependent on their health-state and their level of toxicity. For instance, upon entering a health state a patient may decide to abstain from therapy until their toxicity level falls below a certain threshold. With this extension we are able to jointly consider optimal timing and duration of cancer therapy.

While this model is no longer analytically tractable, we provide the methods for a numerical approximation of evaluating these more general treatment strategies. Patients therefore may select a treatment strategy that maximizes their approximate discounted expected QALYs when affected by toxicity. We also provide an analytical result to calibrate the myopically optimal time of delay of one more round of therapy. If the cure rate decreases in the number of rounds sharply, myopically optimizing the delay of the next round without taking into account any possible future rounds of therapy gives very similar results as approximations of the globally optimal solution.

The paper proceeds as follows. In Section 2 we introduce our base model focusing on the optimal duration of therapy and present a numerical example on the effects of overtreatment. Section 3 adds toxicity to the base model and presents results on the optimal duration and timing of therapy with two numerical examples. Section 4 provides concluding discussions. All proofs are provided in the appendix.

## 2 State-dependent payoffs

We assume that the patient has a solid tissue detectable tumor without specifying the exact kind of cancer. The progression of the disease is modeled as a Markov-process in continuous time. The states encode the patient’s quality of life and prognosis-relevant data, while the transition rates describe their prognosis and depend upon the patient’s chosen treatment strategy. Our state space is given by the set 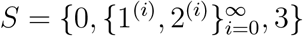. The states are interpreted as follows:

- 0: Healthy, cancer free state.
- 1^(*i*)^: Undetectable cancer after *i* rounds of therapy.
- 2^(*i*)^: Detectable cancer after *i* rounds of therapy. The patient chooses whether to take another round of therapy.
- 3: Death of the patient.

Without therapy, the natural progression of the disease is the following: State 1^(*i*)^ leads eventually to state 2^(*i*)^, state 2^(*i*)^ leads to state 3, an absorbing state. The healthy absorbing state 0 may only be reached by therapy. The patient or the treating physician cannot distinguish the states with undetectable cancer, 0 and 1^(*i*)^, and hence therapy can only be chosen and received while the patient is in a state 2^(*i*)^.

By taking therapy, the patient changes the progression of the disease. If the patient chooses to receive therapy in state 2^(*i*)^ he or she may transition to any one of the four states 0, 1^(*i*+1)^, 2^(*i*+1)^, or 3. Transitioning to 0 and 1^(*i*+1)^ represent therapy success and partial therapy success, respectively, transitioning to 3 and 2^(*i*+1)^ represent therapy failure and partial therapy failure, respectively. The increase of the index from *i* to *i*+1 represents that one round of therapy affects the efficacy of the next one, and thus, although the progression rules remain the same after the *i*th round of therapy as before, the exact transition probabilities may be different. This feature of the game represents, among other factors, the build-up of resistance within the tumor: e.g. reaching state 0 as the result of the *i* + 1th round may be less likely than by the *i*th round.

A *treatment strategy* is characterized by a function 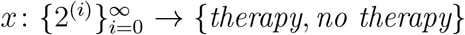. In words, for every state in which the patient has the option to choose, he or she must specify whether or not to take therapy. As a state 2^(*i*+1)^ can only be reached if the patient chooses to receive therapy in state 2^(*i*)^, we restrict attention to treatment strategies such that for every *i* ≥ 0 with *x*(2^(*i*)^) = *no therapy* we have *x*(2^(*i*+1)^) = *no therapy*. We therefore associate a treatment strategy with the maximum number of rounds of therapy the patient chooses to take: *x*_*i*_ means that the patient takes at most *i* rounds of therapy. In strategy *x*_0_ the patient goes without therapy entirely, in strategy *x*_∞_ he or she always opts for therapy when given the choice until reaching an absorbing state.

Time is continuous. We assume that the states encode all progression-relevant information to the disease. Hence the process, conditional on the treatment strategy, is Markovian. The transition rates by which the patient moves between the states are as follows:

1. 1^(*i*)^ → 2^(*i*)^ at rate *δ* _*i*_,
2. if *x*(2^(*i*)^) =*no therapy*, then 2^(*i*)^→ 3 at rate *ω* _*i*_
3. if *x*(2^(*i*)^) = *therapy*,then
  a. 2^(*i*)^→ 0 at rate *λ*_*i*_
  b. 2^(*i*)^→ 1^(*i*+1)^ at rate *β*_*i*_
  c. 2^(*i*)^ → 2^(*i*+1)^ at rate *γ*_*i*_
  d. 2^(*i*)^ → 3 at rate *µ*_*i*_

We introduce the notation *α*_*i*_ = *λ*_*i*_ + *β*_*i*_ + *γ*_*i*_ + *µ*_*i*_. The model’s states and possible transitions are summarized by Figure 1.^2^

**Figure 1:**
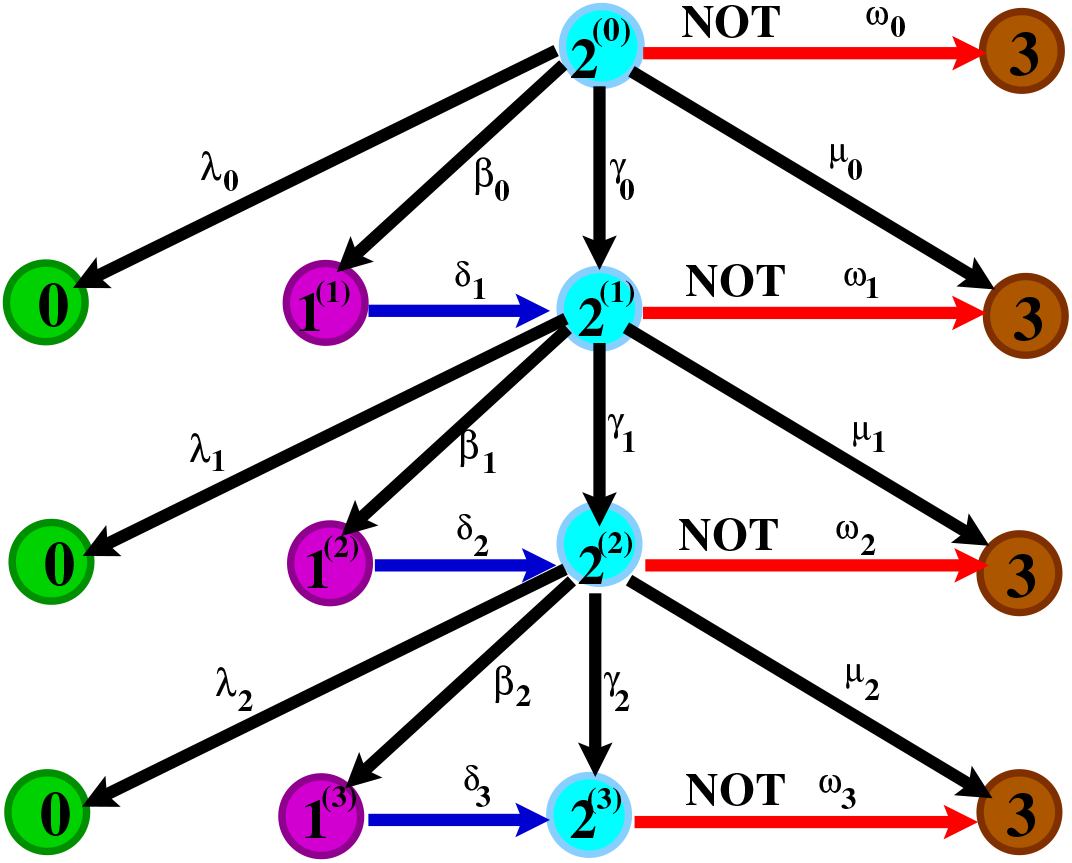
Schematic of transitions of the first 3 rounds of therapy. Each 0 node and each 3 node on the figure represent one absorbing state, the figure shows multiple copies for better visibility. If the patient opts for therapy, he or she progresses to one of the states in the next round. Otherwise, by choosing the **no t**herapy option, he or she eventually progresses to state 3.

Spending time in each health state provides payoffs to the patient measured in QALYs. For this section we assume that this is independent of the chosen treatment strategy. For 0 ≤ *v* ≤ *u* ≤ 1 the function *u* : *S* → [0, 1] given by

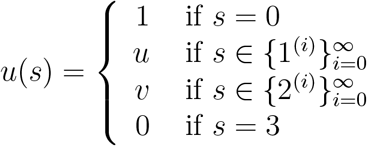

is called the patient’s *instantaneous payoff function*.

Upon selecting the treatment strategy *x*_*i*_, the patient’s progression through the states is a stochastic (Markovian) process. A realization of the patient’s progression is called a play, described by a class of functions *s* : [0, ∞) × *X* → *S*. The value *s*(*t, x*_*i*_), denotes the patient’s state at time *t* ∈ [0, ∞) under treatment strategy *x*_*i*_. Given strategy *x*_*i*_, realization *s*(·, *x*_*i*_) and *j* ≤ *i* let *t*_*j*_(*s*(·, *x*_*i*_)) denote the time that the patient receives the *j*th round of therapy. Whenever it does not cause confusion suppress the argument and write only *t*_*j*_ to denote the time of round *j*.

Taking therapy is costly. Each time the patient accepts therapy he or she instantly incurs a cost *c*. This may represent the monetary cost to pay for one round, lost income, or temporary discomfort caused by the therapy.

We assume that the patient has a preference for earlier rewards, modeled via exponential discounting with discount factor *ρ >* 0.

Given a strategy *x*_*i*_ and realization *s*(·, *x*_*i*_), the patient’s payoffs are given as

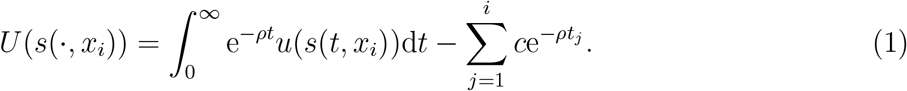

Due to *ρ >* 0, *U* (*s*(·, *x*_*i*_)) is finite for every realization if *i* is finite and for almost every realization if *i* is infinite.

For *j* ≤ *i* let

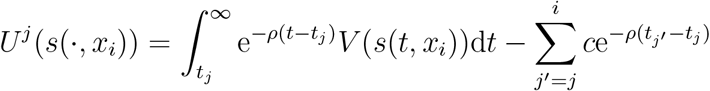

denote the future payoffs of a patient who evaluates their prospect starting from state 2^(*j*)^ (and therefore, starts discounting at *t*_*j*_).

The patient chooses *x*_*i*_ to maximize their expected payoffs given by

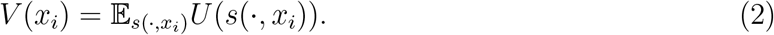

As before, for *j* ≤ *i* we let

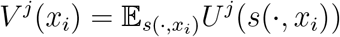

denote the expected payoff of a patient who starts evaluating their prospects from state 2^(*j*)^.

With this we can present this section’s main result on the evaluation of a treatment strategy.

**Proposition 2.1**

(Recursive evaluation) *For a fixed treatment strategy x*_*i*_ *with i >* 0, *the expected future payoffs in round j < i is given as follows:*

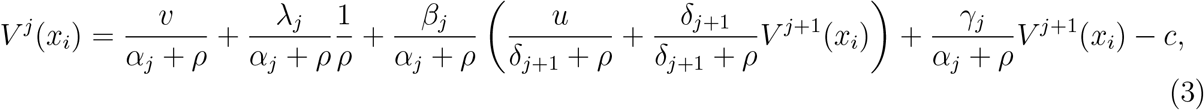

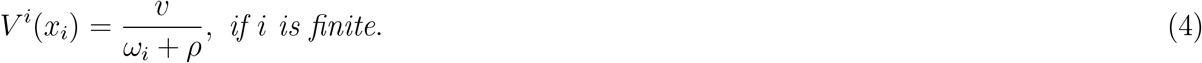

Proposition 2.1 allows for the evaluation of the patient’s payoffs in any state for any finite treatment through a linear recursive system. The right hand side of (3)’s five components are the discounted expected payoff the patient collects in state 2^(*j*)^ before transitioning to any other state; discounted expected value of reaching state 0; discounted expected value of transitioning to state 1^(*j*+1)^, followed by a transition into state 2^(*j*+1)^; discounted expected value of a direct transition to state 2^(*j*+1)^; and the instantaneous cost of the treatment. In (4), as there are no further rounds of therapy, the patient will progress to state 3, thus the right hand side contains only the discounted expected value the patient collects in state 2^(*i*)^ before doing so. In the appendix we calculate each component and formally prove this result.

If for two treatment strategies, *x*_*i*_, *x*_*j*_, we have *V* (*x*_*i*_) ≥ *V* (*x*_*j*_) (*V* (*x*_*i*_) *> V* (*x*_*j*_)) we say that the patient *(strictly) prefers i* to *j* and denote it by *x*_*i*_ ≿ *x*_*j*_ (*x*_*i*_ ≻ *x*_*j*_). We say that *x*_*i*_ is *optimal* if *x*_*i*_ ≿ *x*_*j*_ for every *j*.

Proposition 2.1 allows for optimal treatment strategies to be derived efficiently even though, due to the time-heterogeneity of the transition rates, a closed form of (3) cannot be given. However, (3)-(4) can be transformed to a very simple comparison between two “successive” strategies *x*_*i*_ and *x*_*i*+1_, giving a myopic stopping condition of therapy. This is shown in the next proposition.

**Proposition 2.2**

(Myopic stopping condition) *For a finite i we have x*_*i*_ ≿ *x*_*i*+1_ *if and only if*

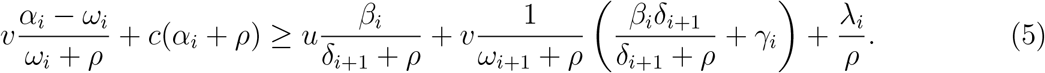

The interpretation is as follows. The advantage of stopping treatment (left-hand-side of (5)) comes from the extra value from spending time in 2^(*i*)^ (*v* term, possibly negative if no therapy results in spending less time in expectation), plus the saved cost of treatment normalized. The advantage of getting another round of treatment (right-hand-side of (5)) comes from the value of spending time in 1^(*i*+1)^ (*u* term), the value of spending time in 2^(*i*+1)^, either indirectly through 1^(*i*+1)^ or by a direct transition (*v* terms), and the value of possibly reaching 0.

Proposition 2.2 can be used to determine if, at any point, stopping therapy immediately is better than continuing once more with the intention to not take any further rounds afterwards. A sequence of such successive comparisons allow for a “local” optimization of the treatment strategy, but, in the general case, not for “global” optimization, for instance, stopping treatment may be better than continuing for one more round, but worse than continuing for two more rounds.

Under certain plausible, or at least possible, monotonicity conditions, however, such local comparisons may give rise to a global optimum, e.g. if continuing for one more round is always better than stopping, then treatment should never be stopped. The last result of this section provides sufficient monotonicity conditions under which the optimal treatment strategy can be calculated by local comparisons.

Take the following homogeneity/monotonicity conditions:

- (H1):*u* = *v*= 1,
- (H2): *δ* _*i*_ = *δ, ω* _*i*_ = *ω*,
- (M1):*M*(*i*) ≤ *M*(*i* + 1),
- (M2):*M*(*i*) ≥ *M*(*i* + 1),

for all *i* ∈ ℕ, and

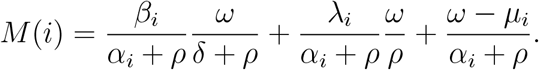

The value *M* (*i*) is a measure of the advantage of taking therapy at state 2^(*i*)^; it is a weighted sum of the progression rates corresponding to at least partial therapy success (i.e. leading to states 1^(*i*+1)^ and 0) and the difference between the death rate without and with therapy.

The first condition is on the patient’s preferences: under (H1) the patient maximizes discounted life expectancy, as time spent in any state other than 3 has the same value. Regarding the transition probabilities: (H2) introduces time homogeneity of the transition probabilities not involving therapy, the rate by which undetectable cancer returns and presents as detectable cancer, and the rate by which untreated patients progress; under monotonicity condition (M1) the patient is *improving* under continuous therapy, the measure of the advantage of taking therapy is increasing in the number of rounds, while under (M2) the reverse holds, the patient’s prognosis is *regressing* under continuous therapy as the measure of the advantage of taking therapy decreases in the number of rounds.

### Proposition 2.3

(Myopic optimization) *Assume (H1) and (H2)*.

1. *Under (M1) there exists an i*′ ∈ ℕ ∪ {∞} *such that for every j < i* ≤ *i*′ *we have x*_*i*_ ≺ *x*_*j*_ *and for every i > j* ≥ *i*′ *we have x*_*i*_ ≿ *x*_*j*_.
2. *Under (M2) there exists an i*′ ∈ ℕ ∪ {∞} *such that for every j < i* ≤ *i*′ *we have x*_*i*_ ≻ *x*_*j*_ *and for every i > j* ≥ *i*′ *we have x*_*i*_ ; ≾ *x*_*j*_.

In the appendix we show Proposition 2.3 by relying on the successive comparisons of Proposition 2.2. Under the first set of conditions, *V* (*x*_*i*_) is quasi-convex in *i*, while under the second it is quasi-concave. In either case we can determine the optimal treatment strategy, as reported in the next corollary.

### Corollary 2.4

(Myopic optimization). *Assumer (H1) and (H2)*.

1. *Under (M1), if V* (*x*_0_) *> V* (*x*_∞_), *then x*_0_ *is the only optimal treatment strategy, if V* (*x*_0_) *< V* (*x*_∞_), *then x*_∞_ *is the only optimal treatment strategy, in case of equality both are optimal*.
2. *Under (M2) x*_*i*′_ *is the only optimal treatment strategy*.

In the first statement, to approximate *V* (*x*_∞_) one can take a sufficiently high *i* and evaluate *V* (*x*_*i*_) through (3)-(4). As we have *ρ >* 0, any level of approximation can be achieved. In the second statement, finding the optimal *i*′ is possible through a sequence of successive comparisons: as long as continuing with one more round of therapy is better than stopping immediately, the patient can continue. Thus, a myopic treatment plan is able to identify the globally optimal treatment strategy.

Naturally, Corollary 2.4 is directly applicable only if the homogeneity and monotonicity conditions (H1), (H2), and one of (M1) and (M2) hold. We argue, however, that its implication is broader. The strategy *x*_∞_ is found to be optimal under an optimistic set of assumptions, specifically that more rounds of therapy improve a measure of the patient’s chances of recovery. Condition (M2), on the other hand is satisfied under more pessimistic parameter settings, and is a closer fit with models of tumor resistance. As a round of therapy is affecting only sensitive cells, further rounds are likely to provide diminishing returns. Under this condition, there exists an interior optimal treatment strategy and any further treatment is to the detriment of the patient.

### Example 2.5

(Overtreatment). In the remainder of this section we simulate the effects of overtreatment and calculate the value lost. To reduce the number of moving parts we introduce a final homogeneity condition, (H3): *β*_*i*_ = *β, γ*_*i*_ = *γ, µ*_*i*_ = *µ*. Under (H1), (H2), and (H3), only the rate of reaching state 0 by therapy, *λ*_*i*_, depends on the number of rounds taken by the patient. We take *λ*_*i*_ = *λ*^(*i*+1)^ for some initial value *λ*. The time-homogeneous parameters of this simulation are shown in Table 1. The effect of varying *λ* and *c* is shown in Figure 2. As expected, the optimal duration of therapy increases with *λ* and decreases with *c*. For *λ* = 0.4 and *c* = 3 the parameters satisfy (M2) and the unique optimal strategy is *x*_2_. Expected values of treatment strategies *x*_0_ through *x*_7_, and the percentage of these compared to the payoffs of a healthy individual are reported in Table 2.^3^

**Table 1:**
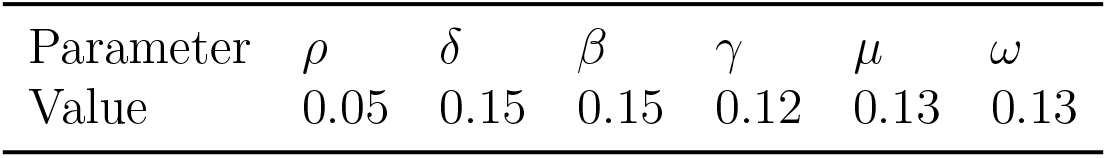
The calibration of Example 2.5. After *ρ* was fixed, the other transition parameters were randomized values between 0.1 and 0.2, keeping *ω* = *µ*, under which a decreasing *M* (*i*) is guaranteed as long as *λ*_*i*_ is also decreasing.

**Table 2:**
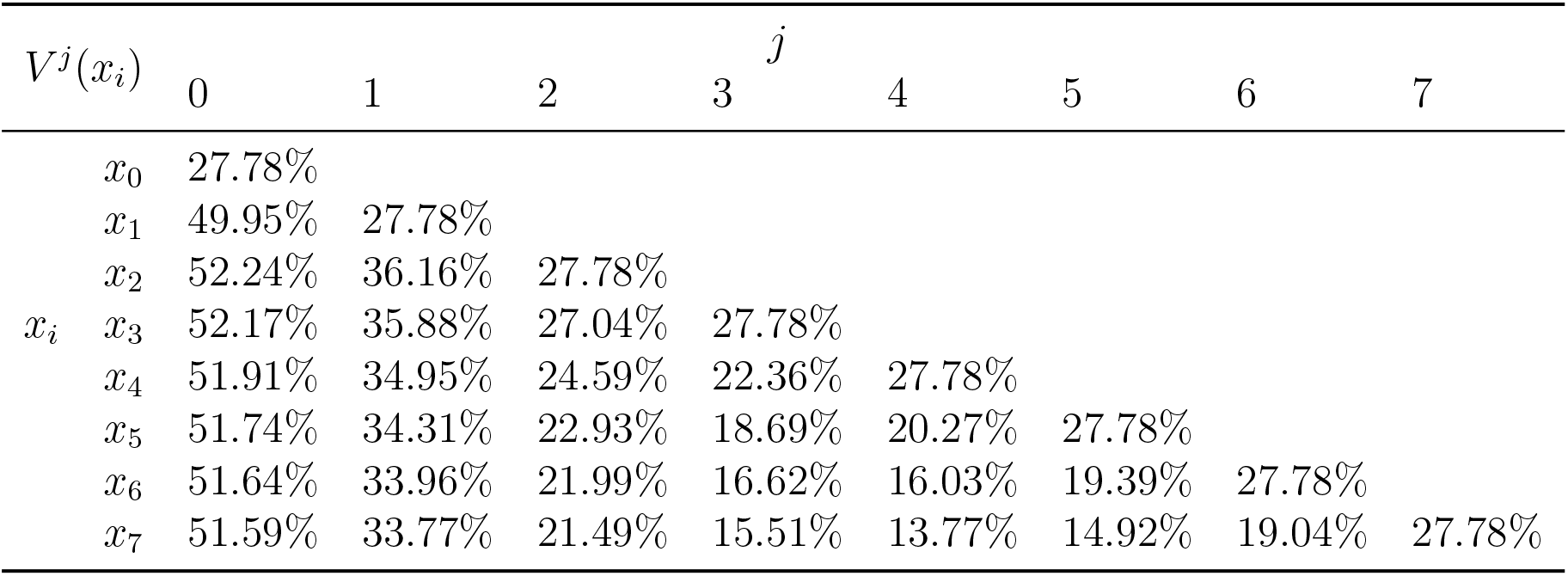
Expected values of treatment strategies *x*_0_ to *x*_7_ evaluated in different time periods relative to a healthy individual’s total payoffs with *λ* = 0.4 and *c* = 3. Taking 2 rounds is optimal, but further rounds diminish the present value (period 0) payoffs only marginally. Patients under continuous therapy who reach round 3 and beyond, if overtreated, have significantly lower prospects than patients who stop therapy.

**Figure 2:**
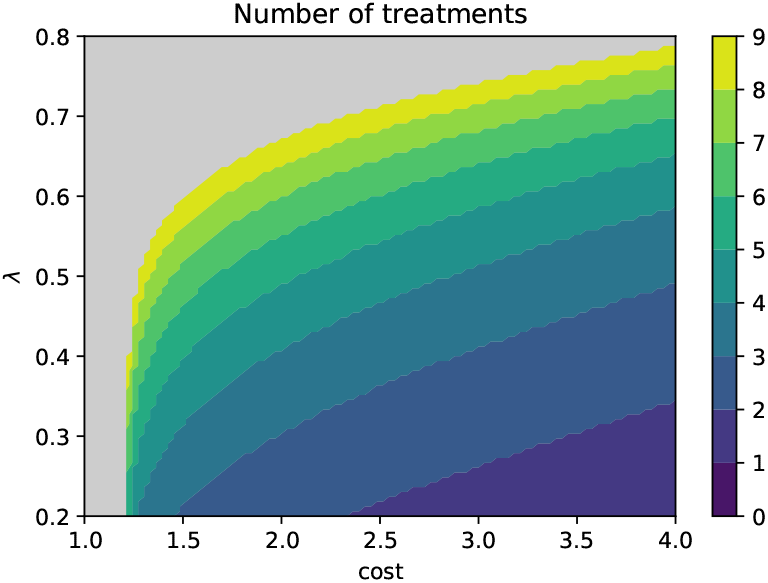
Optimal number of treatment rounds in the cost-based model for parameter values shown in table 1. Gray areas show the regions in which ‘always treat’ is optimal.

As shown by Table 2, any treatment strategy with therapy is better than *x*_0_ with slight variation in the present values, and *x*_2_ being the optimal strategy. However, as *λ*_*i*_ declines sharply, most patients who do not reach state 0 in the first two rounds lose the opportunity to do so in future rounds (Table 3).^4^ For such patients, the cost of future rounds is higher than the present value of the gains of postponing progression to state 3. If the standard of care is continuing therapy indefinitely, patients who reach beyond state 2^(3)^ are overtreated and incur significant payoff losses. Patients reaching round 3 lose 6.29% points under strategy *x*_7_ when compared to the then-optimal *x*_3_, patients who reach round 4 lose 12.27%, while patients who reach round 5 lose the most at 14.01% of a healthy person’s lifetime payoffs. Treatment strategies *x*_1_ through *x*_7_ all provide very similar ex-ante evaluations despite the staggering payoff losses described above. This is due to two reasons: (1) the losses affect a minority of the population (only 9.47% of the cohort is in a non-absorbing state after the third treatment, 6.01% after the fourth, 3.95% after the fifth), (2) the losses occur with a time delay starting in round 3, hence the differences are in the discounted future expected payoffs. Hence, the losses that occur due to overtreatment are obscured, delayed, and concentrated on a minority of patients making policy change to move away from the ‘always treat’ strategy in the standard of care very difficult.

**Table 3:**
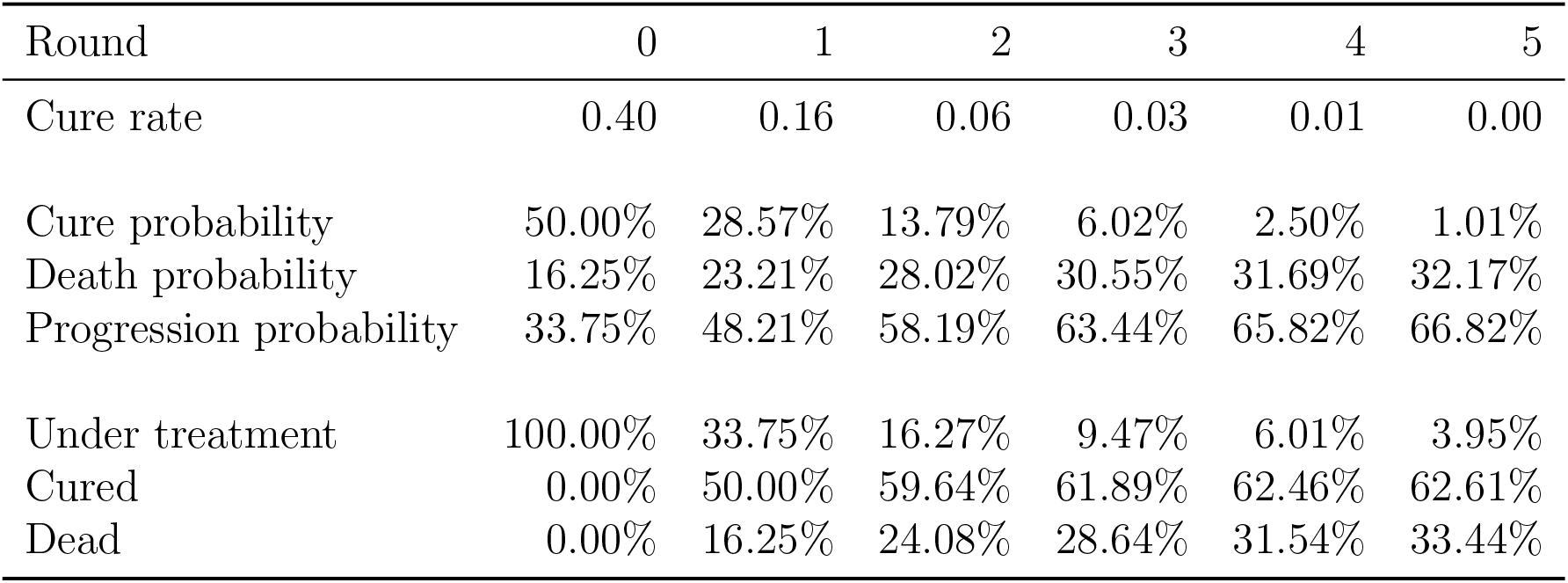
A simulated cohort’s survival statistics under ‘always treat’ with *λ* = 0.4 up to 6 rounds.

It should also be noted that, while in our model and simulation, overtreatment is costly in payoff terms, more patients are cured under treatment strategies with more treatments: *x*_2_ ends with 59.64% of patients cured, while under *x*_7_ this percentage is 62.66%. Furthermore, a payoff-maximizing patient who stops after two rounds refuses the third round despite its cure percentage of 13.79%, showcasing how the objectives of oncologists and patients might differ and lead to highly different choices of treatment strategy.

## 3 Toxicity-dependent payoffs

In Section 2, we modeled the patient’s main restriction for taking therapy by its cost without explicitly mentioning the type of cost element. Such an approach produces a simple and efficient model. Yet, to acquire a deeper understanding of the patient’s choices we need a model that disentangles the material cost elements from those that directly affect the patient’s quality of life.

Cancer patients often incur lifestyle limitations for extended periods. Some of these arise from the disease, while some are due to the side effects of cytotoxic therapies. These side effects, rather than producing a one-time reduction to the patient’s well-being at the time of receiving therapy as we had modeled previously, accumulate and carry over from previous treatment rounds. In particular when a round of therapy is unsuccessful in curing the disease, its lasting effects can influence the decision to take the next round. Instead of the instantaneous reduction, it is useful to model these persisting negative effects as the patient’s ‘rolling stock’ of negative QALYs. We refer to this stock as the patient’s *toxicity*, which increases instantaneously when the patient takes therapy, and decreases over time.

By doing so the scope of our model is also expanded. Previously, the patient’s only decision was in the number of rounds of therapy. Immediately upon entry to a state 2^(*i*)^ the patient chose whether or not to undergo therapy, and the system’s transition rates changed only when the patient entered a new state. In cancer therapy, however, patients also decide on the timing of receiving the next round of therapy. It may be that the patient enters a decision state, spends some time there and risks the *no therapy* rate of progression for some time before deciding to take therapy, after which the *therapy* transition rules apply. In Section 3’s model this behavior is suboptimal as waiting offers no advantage to the patient. However, there are health- and quality of life-related effects of cancer therapy by which delaying the next round is rationally motivated. Modeling the patient’s cost of therapy as a stock allows us to capture these motivations and find the optimal time of delay.

The added ingredient of our model, toxicity, is modeled as follows: Let *i*(*t*) denote the number of rounds of therapy taken up to time *t*. For *z*_0_, 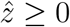 and *ζ >* 0 we define

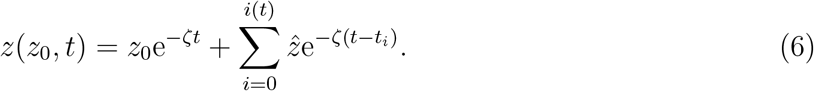

The value *z*(*z*_0_, *t*) is called the patient’s toxicity level, a negative payoff component. Each round of therapy adds a fixed amount 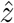 to the patient’s toxicity. Its starting level is denoted by *z*_0_ and it depreciates exponentially with a constant rate *ζ*.

As toxicity is an important component of the patient’s well-being, it becomes a concern for designing treatment strategies. The patient’s choice on the future rounds of therapy is thus contingent on their current level of toxicity. Furthermore, we allow patients to take treatment holidays with the length of holidays also contingent upon the current level of toxicity. Upon entering a state 2^(*i*)^, instead of a binary choice whether to take therapy or not, the patient chooses a time of delay. By delaying for a time 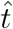, the patient obeys the progression rule as if the *no therapy* choice was taken, i.e. moves to state 3 at rate *ω*_*i*_. If the patient does not progress during this time, then he or she thereafter moves through the game tree in accordance with the *therapy* choice, i.e. moves to state 0, 1^(*i*+1)^, 2^(*i*+1)^, and 3 at rates *λ*_*i*_, *β*_*i*_, *γ*_*i*_, and *µ*_*i*_, respectively.

Formally, the patient’s strategy is now described by a function 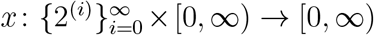. For round *i* and toxicity level *z* the value *x*(*i, z*) is the amount of time the patient waits in state 2^(*i*)^ before administering the next round of therapy. If this value is 0, the next round is administered immediately, if it is infinity, then the patient does not take the *i*+1th round. For consistency, we restrict attention to strategies such that if for some *i* we have *x*(*i, z*) = ∞ for every *z*, then for every *j > i* and every *z*′ we have *x*(*j, z*′) =∞ as well, meaning that if the patient chooses never to take round *i*, all subsequent rounds’ delays are also infinity. We call a treatment strategy *finite* if there exists *i* such that *x*(*i, z*) =∞ for every *z*, i.e. the patient stops therapy after a finite amount of rounds.

The patient’s *instantaneous payoff function when affected by toxicity, u* : *S* × [0, ∞) → ℝ, is given as

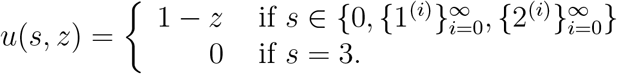

In words, the patient collects a payoff of 1 in any health state other than 3, minus the amount of toxicity he or she currently has. In state 3, the patient collects a payoff of zero. We therefore replace the state-dependent quality-of-life-terms under therapy of our base model, *u* and *v*, with the toxicity-adjusted quality of life, 1 − *z*.

Given a treatment strategy *x*, state-realization *s*(·, *x*) and initial toxicity level *z*_0_, the patient’s *payoff when affected by toxicity* is given by

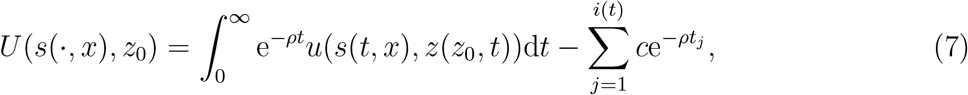

where, as before *t*_*j*_ denotes the time of administering the *j*th round of therapy. Due to *ρ >* 0, *U* (*s*(·, *x*), *z*(·, *x*)) is finite for every realization in every finite strategy and almost every realization for every strategy. We define a patient’s prospects starting in a general state 2^(*i*)^, conditional on the fact that their current toxicity level equals *z*_*i*_ as

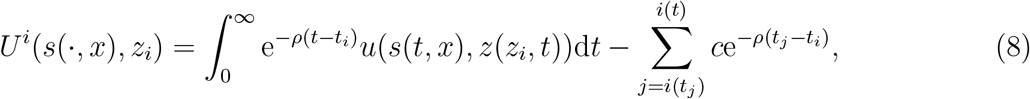

Given *z*_0_, the patient chooses *x* to maximize their *discounted expected payoff* :

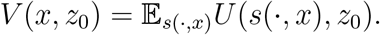

A patient’s who begins the game in state 2^(*i*)^ with toxicity level *z*_*i*_ has prospects given as

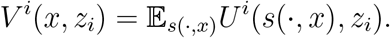

In the following proposition we establish how to evaluate a treatment strategy of a patient affected by toxicity.

**Proposition 3.1**

(Evaluation of treatment strategies under toxicity). *At stage* 2^(*i*)^, *for a treatment strategy x, with starting toxicity level z*_*i*_ *and where the patient waits time* 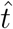 *before taking round i (i*.*e*. 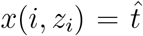*), the patient’s discounted expected payoff is given by the following recursive formula:*

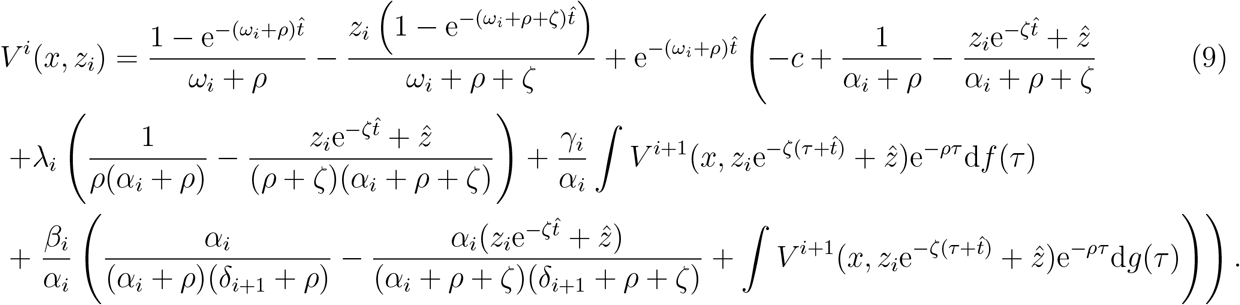

*with probability measures*

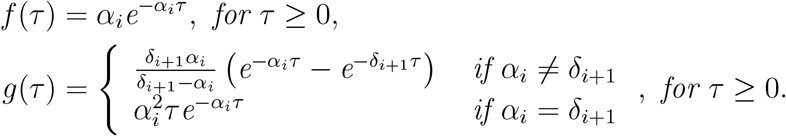

Proposition 3.1 shows the relationship between the payoffs of treatment strategies in successive rounds. The first component is the expected payoff the patient collects while waiting for the next round of therapy. The second component is the sum of three parts: the expected payoff of transitioning to state 0, the expected payoff of a direct transition to state 2^(*i*+1)^, and the expected payoff of a transition to state 2^(*i*+1)^ via state 1^(*i*+1)^.

It is clear that the rolling-stock model of toxicity provides significantly less analytic tractability than the instantaneous cost model of Section 2. This is most apparent by a comparison between Proposition 2.1’s and Proposition 3.1’s respective recursive formulae. While the former shows a simple linear dependence of successive payoff states, the latter necessitates numerical methods of approximation. At the end of this section we examine a numerical example relying on such methods.

In special cases the toxicity model also provides analytically tractable results. Namely, a myopic calibration of the next round’s delay, with the assumption that no further rounds will be taken, is possible. In the next lemma we thus turn to evaluating finite treatment strategies close to the end of treatment. These provide optimal stopping conditions for myopic treatment strategies, and provide insights into a global optimization of treatment strategies. For *i* ∈ ℕ let *X*_*i*_ = {*x* : *x*(*i, z*) = ∞ for all *z*}. Due to the consistency restriction these sets are nested, i.e. *X*_*i*_ ⊆ *X*_*i*+1_ for every *i*.

Let

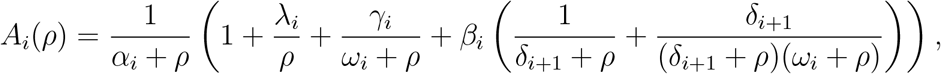

and

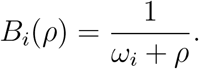

**Lemma 3.2** (Evaluating treatment strategies). *1. For x* ∈ *X*_*i*_

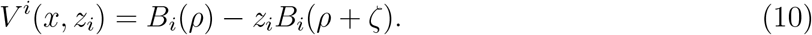

*2. For x* ∈ *X*_*i*+1_ *with x*(*i,z*_*i*_)=0

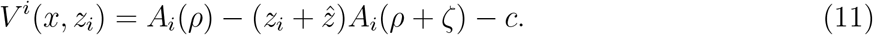

*3. x* ∈ *X*_*i*+1_ *with* 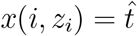

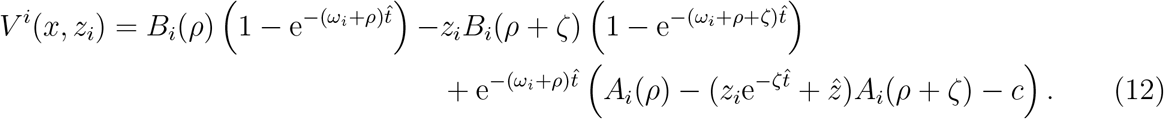

Lemma 3.2 allows us to myopically calibrate the optimal delay before the next round of therapy under the assumption that no further rounds will be taken.

### Proposition 3.3

(Myopic calibration of delay). *Of the strategies with at most i rounds of therapy:*

- *If* 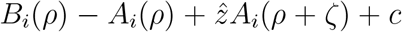 *and B*_*i*_(*ρ* + *ζ*) − *A*_*i*_(*ρ* + *ζ*) *are both negative, then the optimal time to administer the last round of therapy is to wait until the patient’s toxicity level reaches a threshold* 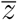 *with*

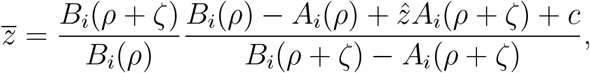

*or, if the patient’s toxicity is below this level, then administer the last round of therapy immediately*.
- *If* 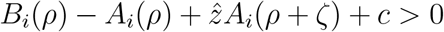 *and B*_*i*_(*ρ* + *ζ*) − *A*_*i*_(*ρ* + *ζ*) *<* 0, *then stopping at the i* − 1*th round is better than continuing with the ith round*.
- *If* 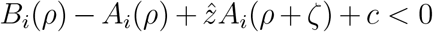 *and B*_*i*_(*ρ* + *ζ*) −*A*_*i*_(*ρ* + *ζ*) *>* 0, *then treatment should be administered immediately*.
- *If* 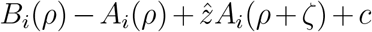 *and B*_*i*_(*ρ* + *ζ*) −*A*_*i*_(*ρ* + *ζ*) *are both positive, then treatment should be administered immediately if the patient’s toxicity is above the threshold z*′ *and never if it is below it, with*

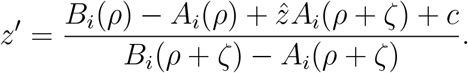

Proposition 3.3 plays a similar role as Section 2‘s Proposition 2.2. It identifies a myopically optimal stopping condition of one round of therapy without an intention of resuming therapy with subsequent rounds. Moreover, it determines the myopically optimal waiting time through analytic methods. Under condition (1) treatment is to be delayed until toxicity is sufficiently diminished, under (2) it is to be canceled no matter the patient’s toxicity level, under (3) it is to be administered immediately no matter the patient’s toxicity level, and finally, under (4) it is to be administered only for patients with high toxicity level. The final point shows a perverse case, resulting from the fact that patients with high negative instantaneous payoffs prefer to immediately receive the next round even when it decreases their life expectancy.

### Example 3.4

We now demonstrate the gains of calibrating the time of delivering the rounds of therapy. As in our previous numerical example, we let *λ*_*i*_ = *λ*^*i*+1^ for an initial value *λ*. Consider the transition parameters shown in Table 4.

**Table 4:**
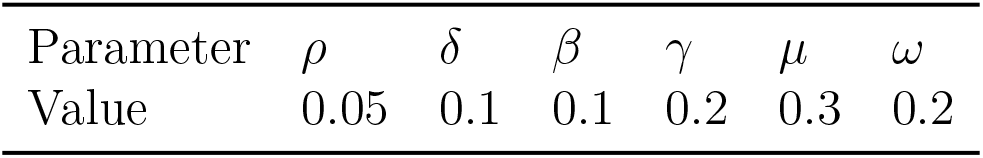
The calibration of Example 3.4.

| Parameter | $\rho$ | $\delta$ | $\beta$ | $\gamma$ | $\mu$ | $\omega$ |
| --- | --- | --- | --- | --- | --- | --- |
| Value | 0.05 | 0.1 | 0.1 | 0.2 | 0.3 | 0.2 |

We first consider the no toxicity case with *λ* = 0.67. Then, as in Example 2.5, (M2) is satisfied. In Table 5, for each treatment strategy *x*_0_ through *x*_8_, we report the cost ranges that produce it as the unique payoff-maximizing strategy.

**Table 5:** Payoff-maximizing treatment strategies for various cost ranges, their corresponding ex-ante payoff ranges relative to a healthy individual, and total cure percentages.

| Cost range | Optimal strategy | Payoff range (% of healthy) | Total cured (%) |
| --- | --- | --- | --- |
| 0.84 – 1.13 | $x_8$ | 64.13% – 62.28% | 65.90% |
| 1.14 – 1.55 | $x_7$ | 62.22% – 59.61% | 65.90% |
| 1.56 – 2.13 | $x_6$ | 59.55% – 55.93% | 65.89% |
| 2.14 – 2.92 | $x_5$ | 55.87% – 50.92% | 65.84% |
| 2.93 – 3.94 | $x_4$ | 50.85% – 44.47% | 65.69% |
| 3.95 – 5.20 | $x_3$ | 44.40% – 36.58% | 65.12% |
| 5.21 – 6.65 | $x_2$ | 36.52% – 27.87% | 62.87% |
| 6.66 – 8.22 | $x_1$ | 27.81% – 20.01% | 52.76% |
| 8.23+ | $x_0$ | 20.00% | 0.00% |

Now consider the case of toxicity. To showcase its effect we set *c* = 0, i.e. the incentive of stopping treatment comes solely from the patient’s decreased quality of life due to toxicity. We take *z*_0_ = 0, 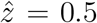, and *ζ* = 0.03. Under these parameters, the “present cost” of one round of therapy due to toxicity is 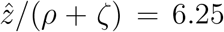. However, this cost is realized in full only by patients with a death rate of zero. Patients in non-absorbing states face a constant death rate of *µ* = *ω* = 0.2 and hence face an “expected present cost” of 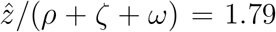. As such, based on Table 5 we can expect at least 2 rounds of therapy and at most 6.

Through Proposition 3.3 we can analytically derive a myopically optimal treatment plan, i.e. the optimal waiting times before each round under the assumption that there will be no further rounds of therapy attempted. As the benefits of further rounds of therapy are declining this will produce increasingly accurate estimates of the globally optimal treatment strategy, starting from that round. In Table 6 we report the threshold levels of toxicity in each round. With *z*_0_ = 0 and 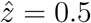 the first two rounds are delivered as soon as possible to the patient as the threshold of round 1 is 0.87, while that of round 2 is 0.76, and the maximum toxicity of the patient after round 1 is 0.5. From round 3 onward, however, the patient may be better off waiting, if their toxicity exceeds the threshold corresponding to round *i* + 1’s at the time of arrival to state 2^(*i*)^.^5^.

**Table 6:** Threshold toxicity levels below which the next round of treatment can be delivered under myopically optimal treatment strategies. Above this level, a payoff-maximizing myopic patient waits until toxicity drops to the threshold level before taking therapy.

| Round | Cure rate | Cure percentage | Threshold toxicity |
| --- | --- | --- | --- |
| 1 | 0.67 | 52.76% | 0.87 |
| 2 | 0.45 | 42.80% | 0.76 |
| 3 | 0.30 | 33.39% | 0.65 |
| 4 | 0.20 | 25.14% | 0.48 |
| 5 | 0.14 | 18.37% | 0.22 |
| 6 | 0.09 | 13.10% | negative |

For a specific case consider a patient in state 2^(2)^, deciding on the delay of the third round This patient has taken two unsuccessful rounds of therapy and their toxicity level increased twice by 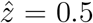, however, in the intermittent times of waiting for the transitions (in states 2^(0)^, 2^(1)^, possibly visiting 1^(1)^ or 1^(2)^ or both as well), the patient’s toxicity level has declined. In our example we set *z*_2_ = 0.73. The patient is facing a cure rate of *λ*_3_ = 0.3. By Table 6, this patient’s payoff is maximized by waiting until the toxicity level reaches 0.65 to take the third round. The patient’s present value, depending on their delay of taking the third round is shown in Figure 3.

**Figure 3:**
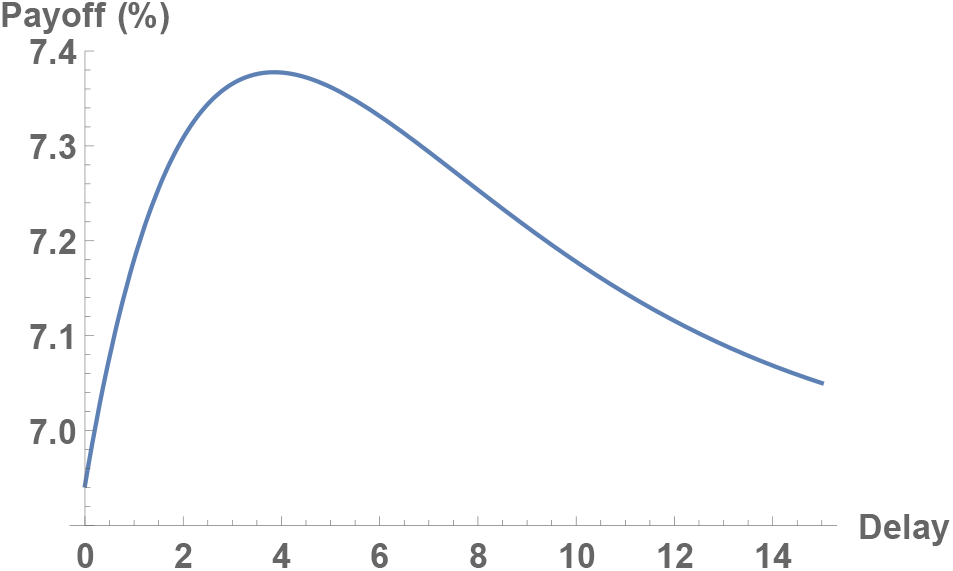
The patient’s payoffs relative to a healthy individual’s after completing two rounds as a function of round 3’s delay with toxicity rate *z*_2_ = 0.73 and facing a cure rate of *λ*_2_ = 0.3. Expected payoffs are maximized at a delay of 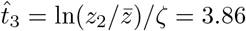

We note that the patient’s decision to delay the third round may seem surprising, considering that the probability of cure is still high (33.39%), and that during the waiting time of 3.86 their probability of death is even higher (e^−3.86*ω*^ = 53.88%). It is clear that such a decision is not supported by practices that maximize probability of cure or survival time. The decision to delay is cast in a more favorable light by considering that receiving the toxicity hit of the third round immediately would yield a quality of life of − 0.23 – even at the threshold toxicity of 0.65 the patient’s quality of life turns temporarily negative. Delaying lowers the “present cost” of therapy enough for a payoff-maximizing patient to take it.

Example 3.4 showcases both the possible benefits of delaying therapy (Figure 3) and a myopically optimal patient’s behavior (Table 6). It also highlights the comparison between the models of Sections 2 and 3. The former prescribes the number of treatment rounds based on the flat one-time cost the patient incurs per round, while the latter prescribes the timing of these rounds. Note, however, that unlike in Section 2, where we were able to derive a condition that ensured that the myopically optimal behavior produces the globally optimal one (Proposition 2.3), there is no analogous result to guarantee that Table 6’s results correspond to the globally optimal behavior in the toxicity model. In the next example, we evaluate the same calibration via a numerical approximation and show that its results are in agreement with the myopically optimal waiting times.

### Example 3.5

Consider the same transition parameters as shown in Table 4. As in Example 3.4, we take *λ* = 0.67, 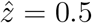 and *ζ* = 0.03 with *z*_0_ = 0. Table 7 reports the expected optimal delays of a maximum of six treatment rounds through a numerical approximation (see the appendix for a summary of the methodology of the approximation).

**Table 7:** Delay times and payoffs of approximate optimal strategies, *x*^*^(*i, z*_*i*_) conditional on starting therapy in round *i* with toxicity level *z*_*i*_. Bold numbers are actionable choices, all other delays are expected values subject to change. A patient progressing through the rounds re-optimizes in each round and tailors their behavior based on the current level of toxicity.

|  |  |  | Round | 0 | 1 | 2 | 3 | 4 | 5 |
| --- | --- | --- | --- | --- | --- | --- | --- | --- | --- |
|  |  |  | Cure rate | 0.67 | 0.45 | 0.30 | 0.20 | 0.14 | 0.09 |
| $i$ | $z_i$ | Payoff | Cure perc. | 52.76% | 42.80% | 33.39% | 25.14% | 18.37% | 13.10% |
| 0 | 0.00 | 42.70% | | **0** | 0 | 0 | 11.27 | 20576 | ∞ |
| 1 | 0.32 | 24.12% | | | **0** | 0 | 13.76 | 17.67 | ∞ |
| 1 | 0.40 | 21.16% | | | **0** | 0.84 | 13.76 | 118.88 | ∞ |
| 1 | 0.48 | 18.28% | | | **0** | 3.68 | 13.91 | 13.77 | ∞ |
| 2 | 0.60 | 11.09% | | | | **0** | 15.44 | 176.77 | ∞ |
| 2 | 0.68 | 8.62% | | | | **1.44** | 16.96 | 22.31 | ∞ |
| 2 | 0.76 | 6.73% | | | | **5.14** | 16.96 | 22.19 | ∞ |
| 2 | 0.84 | 5.13% | | | | **8.48** | 16.96 | 21.88 | ∞ |
| 2 | 0.92 | 3.63% | | | | **11.51** | 16.96 | 24.12 | ∞ |

The interpretation of the prescribed treatment strategy starting at round 0 (first row of Table 7) is as follows: Given the patient’s toxicity level of *z*_0_ = 0, in expectation, the patient is advised to wait time 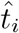 before receiving the *i* + 1th round of therapy. Note that the prescribed waiting times for distant treatment rounds are subject to change. At the onset, they are merely an expected time of optimal delay given the patient’s *expected* progression, on which, based on backwards induction, the optimal time of delay of the first round, 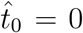, can be calculated. Thus, only this first delay is actionable information. Should the patient reach the next decision node, their toxicity level may be quite different from the expected levels, hence, subsequent decisions need to be taken according to the *realized* toxicity levels.

To illustrate, we report three re-optimized treatment strategies given toxicity levels *z*_1_ = 0.32, 0.40, and 0.48 after round 1 (rows 2 to 4 of Table 6). This large divergence in toxicities is based on the fact that patients who do not respond to the treatment (and thus progress to state 2^(1)^ directly) are expected to have larger toxicity levels than those who do (and thus reach 2^(1)^ indirectly through 1^(1)^), as the latter group’s toxicity depreciates for a longer time.^6^ As shown in the table, these patients are all advised to take round 2 immediately, but their expected delays in future rounds, as well as their expected payoffs, diverge.

Those patients who progress further again need to re-optimize based on their realized levels of toxicity. We approximate optimal treatment strategies for patients who start after round 2 with toxicity levels *z*_2_ = 0.60, 0.68, 0.76, 0.84, and 0.92. At this stage, the prescribed delays before taking round 3 are different, hence the different patients’ payoff-maximizing behavior diverges. The approximate delay times of the next round line up with the myopically optimal ones (retrieved from Proposition 3.3) up to the 3rd decimal point, indicating that the approximate optimal solution and the myopically optimal one agree closely, provided that *λ*_*i*_ is decreasing.

## 4 Concluding discussion

In this paper we built a decision-making tool of cancer therapy. We model the development of the disease as a random, Markovian process, capturing the prognosis-relevant data with four types of health states. This approach unifies the more classical Markovian models of cancer therapy with the novel game theoretic analysis of cancer, adding the element of patient choice to the former, and simplifying cancer’s evolutionary dynamics to a random, Markovian process in the latter. Framing cancer’s strategies such a way allows us to focus on the patient’s choices and rely on classic results of the theory of Markov Decision Processes for the existence of a unique optimal policy: an optimal treatment strategy.

In a model where the patient’s instantaneous payoffs are determined by the type of health state they currently occupy, we provide a simple recursive formula to analytically evaluate the performance of various treatment strategies. Estimating transition rates from cohort data and inputting the parameters reflecting the patient’s preferences allows the patient to choose their preferred therapy duration. Under some monotonicity and homogeneity assumptions, a local and myopic evaluation of the treatment strategies also produces the globally optimal outcome, further simplifying the decision-making progress. In a second model, where the patient’s instantaneous payoffs were determined by their current toxicity levels, the evaluation of treatment strategies is more complicated and requires numerical tools. Nevertheless, optimal duration of therapy and optimal timing of treatment rounds can be estimated. Myopically optimizing the next round’s delay can be performed analytically, and can provide a good approximation to a globally optimal treatment strategy if the cure rate of future rounds decreases sharply.

We raise three discussion points on the modeling choices made in the paper. The first is the decision to include no more than four types of health states. One reason for this is to keep our models tractable. A second reason is that a practical application of a model with more health states requires more cohort data. Given the same amount of cohort data, calibrating a model with more than four health states comes with a loss of statistical power. In the case of large cohorts, collecting patient data of a given cancer type, this may not be a problem. However, in the case of cohorts stratified by age, sex, or by other variables, diluting the data in favor of including a larger number of health states may not be desirable. We further argue that more health states raises classification problems, while the four present in our paper is the lowest number that is needed. In cases where data are abundant and classification unproblematic, our model can be extended to include more state types in a straightforward manner.

Secondly, we raise the issue of personalized medicine. Barring some exceptional circumstances, the transition rates of our model must be calibrated from cohort data. The ability to personalize our model depends on the availability cohort data corresponding to the patient’s stratum. For some cancers and for some strata this cannot be taken as given. In these cases, our models can still serve as useful benchmarks against which the patient and their physician may evaluate their options given the patient’s own characteristics and response. Even when the ability to personalize our model’s transition rates is low, some of our model’s variables such as the patient’s instantaneous payoff parameters and discount rate can be calibrated to match the patient’s preferences and characteristics. When personalization is high, the differences between these patient-specific traits may still mean that two patients belonging to the same demographic will find different treatment strategies optimal.

Thirdly, we address the relationship of the patient’s toxicity level in our second model and the transition rates. In our model, these are mathematically independent in the sense that after a given number of rounds of therapy, progression rates are not affected by toxicity. In practice, toxicity caused by therapy is strongly related to the patient’s prognosis. This mismatch is caused by the fact that our model combines “objective” parameters regarding disease prognosis with “subjective” ones that reflect to the patients’ preferences. Toxicity of therapy is related to both. We therefore use the abstract term toxicity to reflect on the subjective aspect, measuring the patient’s well-being under therapy. Introducing explicit dependence between toxicity and transition would be problematic both for the tractability of the model and in mixing the “objective” concerns with “subjective” ones. For example, two patients may be very similar in their disease progression but may report varying levels of discomfort due to therapy, or vice versa, which may influence their choice of treatment. As the “objective” effects of toxicity, the transition rates, do depend on the number of rounds of therapy, our toxicity measure and the patient’s prognosis are statistically not independent. Hence, our model may produce a good fit even without mathematical dependence between toxicity and transition.

Finally, we reflect on our stated goal, to address the dilemmas arising from the difficulty in finding a suitable measure of success of cancer therapy. Our approach, maximizing the patient’s discounted expected QALYs is rooted in a classic economic approach that treats individuals as rational utility maximizers. As such, we propose it as a good candidate to evaluate cancer therapy in a way that explicitly captures the patients’ well-being. As an additional value, even if such an approach cannot be adopted in oncology formally, a model such as this can help identify and understand points of disagreement between cancer patients and their treating physicians in selecting a treatment strategy.

Our approach shares the drawbacks and criticism of similar decision theory models: (1) QALYs (or payoffs in general) are significantly more difficult to measure than survival, and (2) individual decisions often go against what economists or game theorists describe as “rational”. As an added difficulty, (3) individual decision-making in dynamic situations may be, and is often shown to be, time-inconsistent. Addressing (1) in the cancer context is part of a deeper discussion on the appropriateness of using QALYs. We argue that, while its shortcomings do not make it suitable to replace more convenient measures, such as survival time, considering QALYs in addition to survival time has significant added value. To address (2) and (3) would require a deeper mapping of the individual decision-making process. Methods that are currently used in behavioral economics, psychology, and other decision sciences often use very similar tools as those in strictly “rational” models. Thus, our methodology, as well as its predictions, can serve as useful benchmarks for future research in the decision theory of cancer.

Other important aspects of decision making that have not been captured by our model include the patients’ risk and ambiguity attitudes. Our model assumes perfect information of transition rates and risk neutral patients but both assumptions can be relaxed in a straightforward manner, the former by introducing noise to the transition process, the latter by incorporating the patient’s risk and ambiguity attitude in their (perceived) payoff. Doing so constitutes an important direction of future research. Additionally, while our model is silent on the treating physician’s incentives, the dilemma arising from the physician and the patient’s different objectives can be explicitly captured by a principal-agent problem, of which, our findings represent one side, that of the principal. This direction is also left for future research.

## A Appendix

### Proposition 2.1

We first show the second part of the statement, that is:

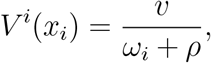

for a finite *i*.

The patient collects a constant stream of instantaneous payoffs *v* while still in state 2^(*i*)^, and 0 after he or she transitions to state 3. Let *τ* denote the time the patient spends in 2^(*i*)^. As *τ* ∼ Exp(*ω*_*i*_), we have

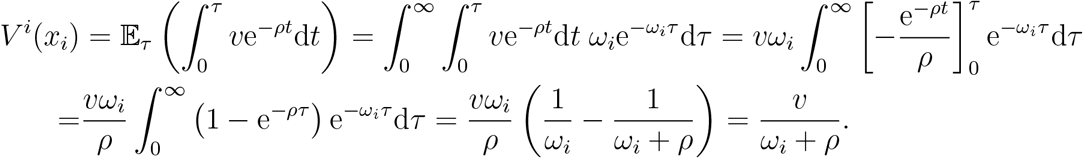

To show the first part we calculate each of the following four components separately: (1) the discounted payoffs collected in state 2^(*j*)^ before transitioning; (2) those collected after transitioning to state 0; (3) those collected after transitioning to state 1^(*j*+1)^, followed by transitioning to state 2^(*j*+1)^; (4) those collected after a direct transition to 2^(*j*+1)^.

Calculating (1) amounts to evaluating

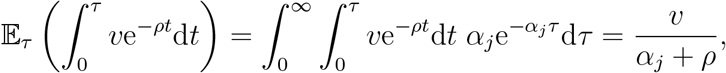

with very similar steps as before, where now we have *τ* ∼ Exp(*α*_*j*_).

To calculate (2) we need to evaluate

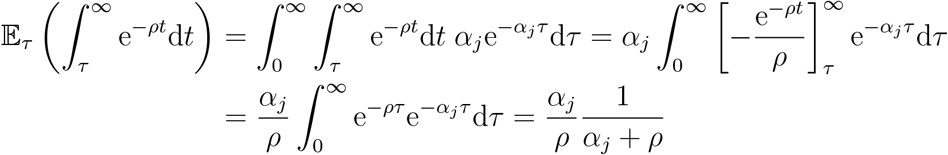

as once more we have *τ* ∼ Exp(*α*_*j*_). Multiplying by *λ*_*j*_*/α*_*j*_, the probability that state 0 is reached, we get

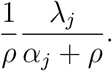

Component (3) has two parts: the payoffs collected while the patient is in state 1^(*j*+1)^, and the payoff he or she collects after transitioning to 2^(*j*+1)^. Taking *τ* ∼ Exp(*α*_*j*_) and *τ*′ ∼ Exp(*δ*_*j*+1_), the former amounts to

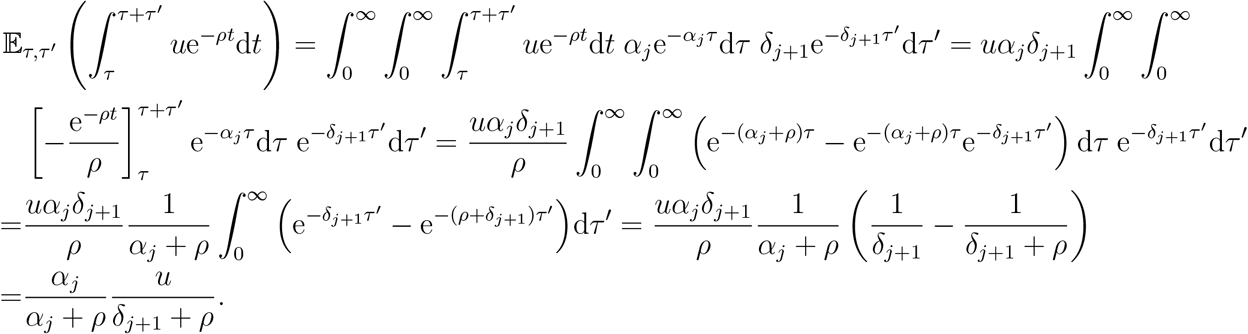

This, multiplied by the probability of reaching state 1^(*j*+1)^, *β*_*j*_*/α*_*j*_ gives

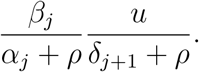

The second part, the payoff the player receives after transitioning to 2^(*j*+1)^ amounts to receiving a payoff of *V* ^*j*+1^(*x*_*i*_) with time delay *τ* + *τ*′, that is, in expectation:

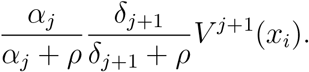

Multiplying by the probability of reaching state 1^(*j*+1)^ (from which reaching state 2^(*j*+1)^ is certain), we get

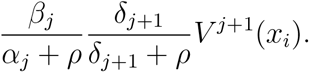

The sum of the two parts gives the third component of (3) as desired.

In component (4), a direct transition to state 2^(*j*+1)^ provides a payoff of *V* ^*j*+1^(*x*_*i*_) with a delay of *τ* with *τ* ∼ Exp(*α*_*j*_), equaling

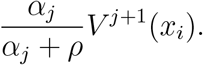

Multiplied by the probability of reaching 2^(*j*+1)^ directly, *γ*_*j*_*/α*_*j*_, we get

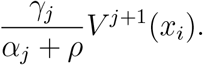

Finally, subtracting the cost of a round of therapy, *c*, incurred immediately, we get the right hand side of (3).

### Proposition 2.2

As the two treatment strategies are identical in the first *i* periods, *V* (*x*_*i*_) ≥*V* (*x*_*i*+1_) if and only if *V* ^*i*^(*x*_*i*_) ≥ *V* ^*i*^(*x*_*i*+1_). By Proposition 2.1 the left hand side amounts to *v/*(*ω*_*i*_ + *ρ*), while the right hand side is

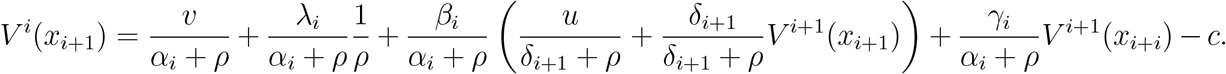

By plugging in *V* ^*i*+1^(*x*_*i*+1_) = *v/*(*ω*_*i*+1_ + *ρ*) we have that *V* ^*i*^(*x*_*i*_) ≥ *V* ^*i*^(*x*_*i*+1_) if and only if

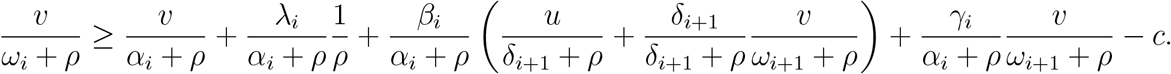

Multiplying by *α*_*i*_ + *ρ* and rearranging produces the inequality stated by the proposition.

### Proposition 2.3

Applying (H1) and (H2) to (5), by Proposition 2.2 we have *x*_*i*_ ≾ *x*_*i*+1_ if and only if

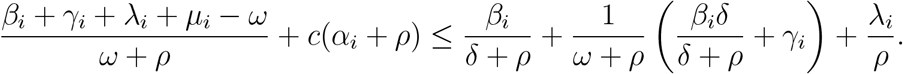

Multiplying by (*ω* + *ρ*)*/*(*α*_*i*_ + *ρ*) and rearranging gives

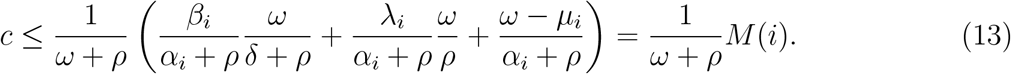

1. Let *i*′ ∈ ℕ be the smallest number such that *x*_*i*′_ ≾ *x*_*i*′ +1_. Then we have *c ≤ M* (*i*′)*/*(*ω* + *ρ*). Under (M1) *M* (*i*) is increasing in *i*, thus every successive treatment strategy with more than *i*′ rounds is better than the one preceding it, hence for every *i > j* ≥ *i*′ we have *x*_*j*_ ≾ *x*_*i*_. By the choice of *i*′, for every *j* ≤ *i*′ *>* 0 we have then *x*_*j*_ ≺ *x*_*j*−1_, implying that for every *i < j* ≤ *i*′ we have *x*_*j*_ ≺*x*_*i*_.
2. Let *i*′ ∈ ℕ be the smallest number such that *x*_*i*′_ ≿ *x*_*i*′ +1_. Then we have *c ≥ M* (*i*′)*/*(*ω* + *ρ*). Under (M2) *M* (*i*) is decreasing in *i*, thus every successive treatment strategy with more than *i*′ rounds is worse than the one preceding it, hence for every *i > j* ≥ *i*′ we have *x*_*j*_ ≿ *x*_*i*_. By the choice of *i*′, for every *j* ≤ *i*′ *>* 0 we have then *x*_*j*_ *≻ x*_*j*−1_, implying that for every *i < j* ≤ *i*′ we have *x*_*j*_ *≻ x*_*i*_.

### Proposition 3.1

The value is the sum of five values: (1) the payoff received in state 2^(*i*)^ while waiting for the next round of therapy. We calculate the positive part of the payoff (i.e, without toxicity). Take *τ* ∼ Exp(*ω*_*i*_), then

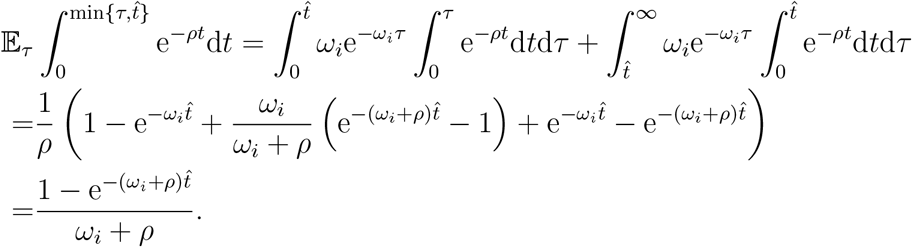

With very similar calculations we may get the negative (toxicity) part of this component:

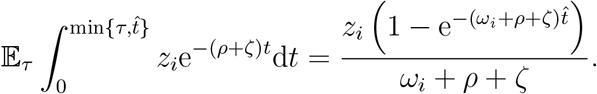

(2), the payoff received in state 2^(*i*)^ after taking therapy but before transitioning to any of the states 0, 1^(*i*+1)^, 2^(*i*+1)^, or 3 as a result. Again, just taking the positive component, with *τ* ∼ Exp(*α*_*i*_) this is

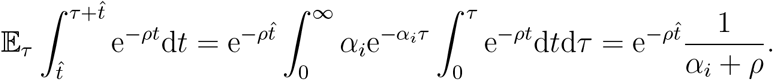

For the toxicity component that the patient started with, we get

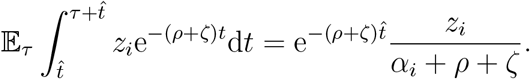

Adding the toxicity caused by therapy 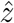 at time time 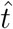 we get

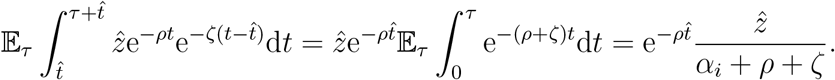

Adding these three and multiplying with the probability of the patient reaching the time to take therapy, 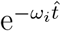 we get

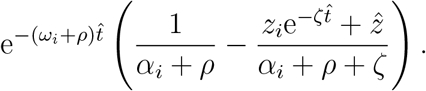

(3), the payoff received upon a transition to state 0. Again, with *τ* ∼ E(*α*_*i*_) this is (positive and negative parts together):

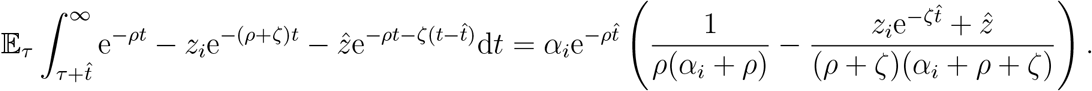

Multiplying with the probability reaching the time to administer round *i*, 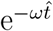, and by the probability of transitioning to state 0 given that the patient receives round *i, λ*_*i*_*/α*_*i*_, we get

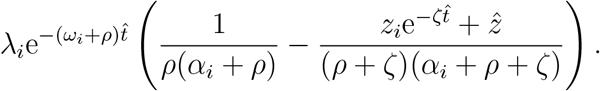

(4), the payoff received upon a transition to state 2^(*i*+1)^. This amounts to the expected present value of *V* ^*i*+1^(*x, z*(*z*_*i*_, *τ*′)) with delay *τ*′ where 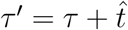 for *τ* ∼ Exp(*α*_*i*_). This equals

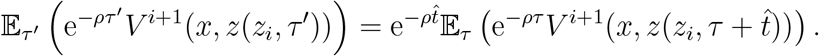

Multiplying by the probability of reaching the time to administer round *i*, and by the probability of transitioning directly to state 2^(*i*+1)^ given that the patient receives round *i, γ*_*i*_*/α*_*i*_ and substituting in 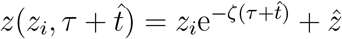 we get

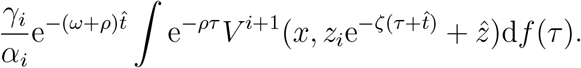

(5), the payoff received upon a transition to state 1^(*i*+1)^ followed by a transition to state 2^(*i*+1)^. With *τ*_1_ ∼ Exp(*α*_*i*_) and *τ*_2_ ∼ Exp(*δ*_*i*+1_), the former amounts to

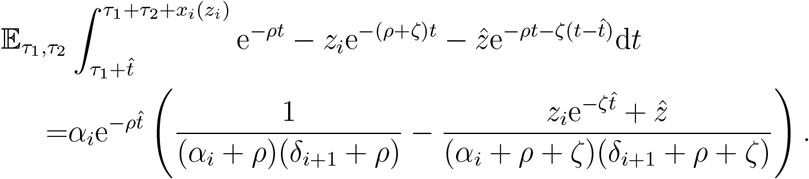

Multiplying by the probability of reaching the time to administer round *i*, and by the probability of transitioning to state 1^(*i*+1)^ from 2^(*i*)^, *β*_*i*_*/α*_*i*_, we get

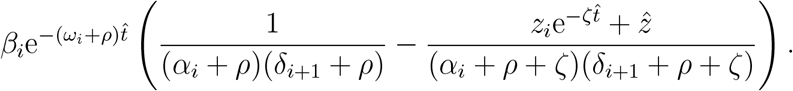

Finally, upon reaching state 2^(*i*+1)^ from 1^(*i*+1^) the patient receives the present expected value of *V* ^*i*+1^(*x, z*(*z*_*i*_, *τ*′)) with a delay of *τ*′ where 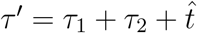. Substituting *τ* = *τ*_1_ + *τ*_2_ we get

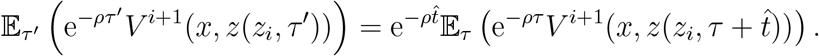

Multiplying by the probability of reaching the time to administer round *i*, and by the probability of transitioning directly to state 1^(*i*+1)^ (from which reaching state 2^(*i*+1)^ is certain) given that the patient receives round *i, β*_*i*_/*α*_*i*_ 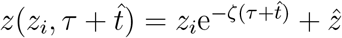 we get

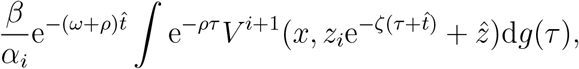

as *g*(·) is the density function of *τ*_1_ + *τ*_2_ by definition.

Summing up components (1) through (5) and adding the cost of one round of therapy, *c* with delay 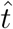 multiplied by the probability of paying it gives the formula stated by the proposition.

### Lemma 3.2

1. (10) is obtained from (9) by setting 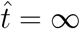.
2. To calculate positive component of the payoff (without toxicity and costs), we substitute 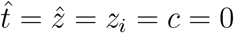 into (9) to obtain

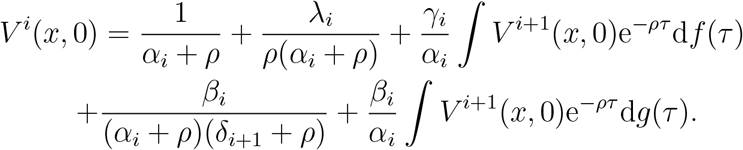 By point 1, we may substitute *V* ^*i*+1^(*x*, 0) = *B*_*i*_(*ρ*). Evaluating the integrals gives

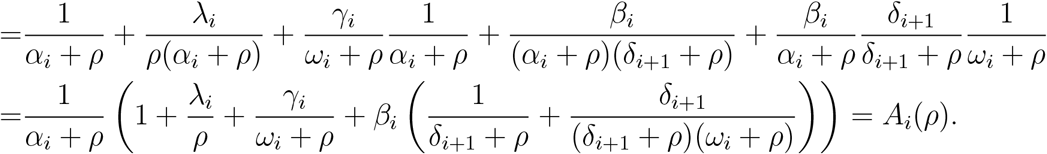 By similar calculations the payoffs from toxicity equal 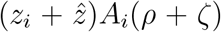, while the cost is a lump-sum −*c*. Adding these together gives (11).
3. Calculating the positive components amounts to substituting 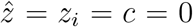 into (9). This yields

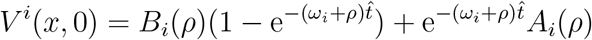

where the second component follows from the calculations of the positive component of 2. The toxicity can be deduced as

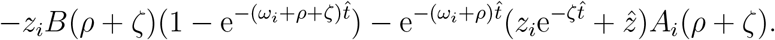

Adding these together with the lump-sum cost − *c*, factoring in the delay and the probability of paying the cost leads to (12) as stated.

### Proposition 3.3

We take a treatment strategy *x* ∈ *X*_*i*+1_ and evaluate it in state 2^(*i*)^ given toxicity level *z*_*i*_. To find the optimal 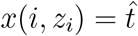 we differentiate *V* ^*i*+1^(*x, z*_*i*_) (deduced from Lemma 3.2) with respect to 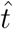 to give

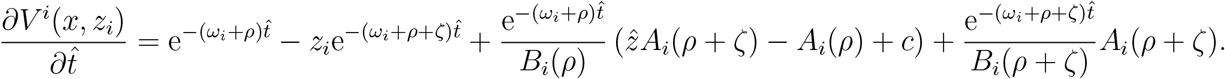

Multiplying by 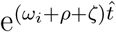 and rearranging, the sign of the derivative is the same as that of

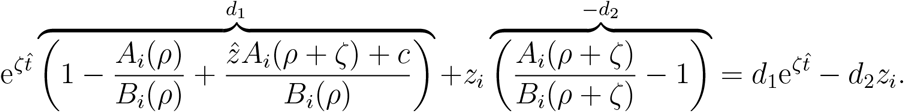

There are four cases: 1. If *d*_1_ and *d*_2_ are both negative, then the derivative equals zero if

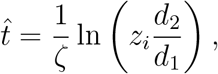

provided that *z*_*i*_ *> d*_1_*/d*_2_. If so, then 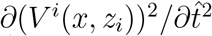 is negative due to *d*_1_ being negative, hence 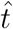 is indeed a maximizer, and 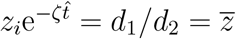, thus the patient waits until toxicity falls to 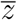. If *z*_*i*_ *< d*_1_*/d*_2_, then the first derivative is always negative, hence taking the next round immediately is optimal.

2. If *d*_1_ *>* 0 and *d*_2_ *<* 0, then the first derivative is positive for all 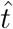, hence 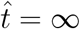 is optimal.

3. If *d*_1_ *<* 0 and *d*_2_ *>* 0, then the first derivative is negative for all 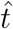, hence 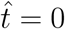 is optimal.

4. If *d*_1_ and *d*_2_ are both positive, then if 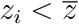, then the first derivative is positive for all 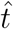, meaning that 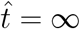 is optimal. If 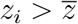, then the first derivative starts negative at 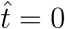, then turns positive and remains positive as 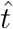 approaches infinity, meaning that either 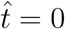 or 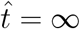 is optimal. Comparing the payoffs, we get that 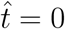 is best if and only if

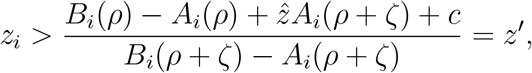

which is a stronger condition than 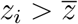.

### Approximation method of Example 3.5

All transition parameters with the exception of the cure rate, *λ*_*i*_, are independent if *i*. We assume a maximum number of treatments, *N*, that is, we set 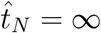.

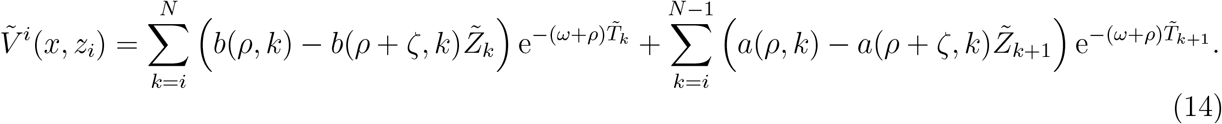

The components in (14) are as follows: We denote by 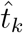 the time of delay before treatment round *k* with 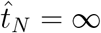. The series *T*_*k*_ denotes the times at which the patient’s toxicity increases as a result of the *k*th round of treatment, which takes place time 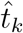 after the patient enters 2^(*k*)^. *T*_*i*_ is taken to be 0, while for *k > i* we have

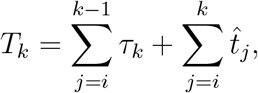

with *τ*_*k*_ being the random variable denoting the length of the *k*th round of therapy from its initiation (i.e. when toxicity increases) to its termination, conditional on the fact that the patient proceeds to state 2^(*k*+1)^.

To get an approximation, we replace *T*_*k*_ in (14) by its expected value, 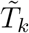, leading to anunbiased estimate of it. Given the patient’s strategy, the waiting times 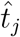 are fixed, while the expected value of *τ*_*k*_ is given by

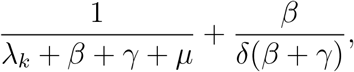

of which the first component is the expected time spent in state 2^(*k*)^ while waiting for the *k*th round to take effect and the second is the expected time spent in state 1^(*k*+1)^, waiting for progression to state 2^(*k*+1)^, leading to 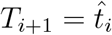

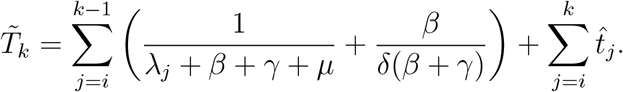

The estimate 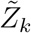 denotes the approximation of the patient’s toxicity at the time of receiving the *k*th therapy, i.e. at time *T*_*k*_. For simplicity and computational ease, we approximate the patient’s toxicity level at the time of entering state 2^(*k*)^ by substituting the expected time into the toxicity equation (6), giving a slightly biased estimate of the patient’s toxicity:^7^

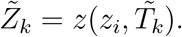

The two major components in (14) are

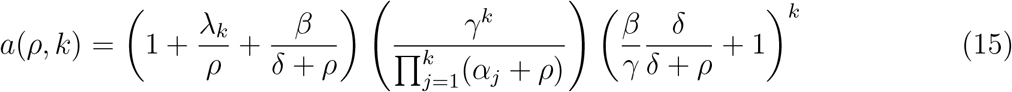

and

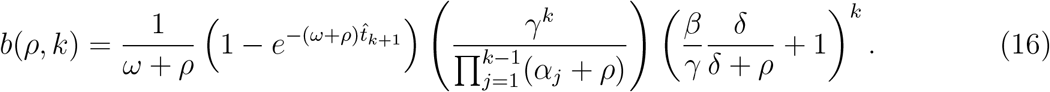

To get a visual intuition in deriving (14), from Figure 1, imagine that we fix the maximum number of treatments at *N*, reducing the model to a finite series of states. We descend *N* layers in the figure, then calculate all the possibilities to arrive at either state 0 or state 3 after at most *N* treatments by simply counting the number of paths. Each new layer can be reached one of two ways, either a direct transition from state 2^(*i*)^ to 2^(*i*+1)^ with rate *γ*, or an indirect one from 2^(*i*)^ to 1^(*i*+1)^ at rate *β*, then from 1^(*i*+1)^ to 2^(*i*+1)^ at rate *δ*.

The approximations of Table 7 are therefore results of numerically maximizing (in Wolfram Mathematica) equations of the form (14), subject to 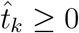, and entering *λ*_*k*_ = *λ*^*k*^ into (15).

## Footnotes

1 In the remainder of the paper we refer to the patient as the sole decision-maker without explicitly mentioning the treating oncologist, tumor board, or any other participants of the decision making process.

2 The connection with the more well-known discrete-time Markov Decision Processes is summarized as follows: In expectation, a patient who does not take therapy at state 2^(*i*)^ spends time 1*/ω*_*i*_ in 2^(*i*)^ before progressing to 3. The transition probability from 2^(*i*)^ to 3 is thus 1 without therapy. Similarly, a patient in 1^(*i*)^ transitions to 2^(*i*)^ with probability 1, spending at expected time of 1*/δ*_*i*_ in 1^(*i*)^. A patient who takes therapy in 2^(*i*)^ spends an expected 1*/α*_*i*_ time in this state before transitioning to one of 0, 1^(*i*+1)^, 2^(*i*+1)^, 3 with probabilities *λ*_*i*_*/α*_*i*_, *β*_*i*_*/α*_*i*_, *γ*_*i*_*/α*_*i*_ and *µ*_*i*_*/α*_*i*_, respectively. The time spent in each state is exponentially distributed with parameter corresponding to the total transition rate out of the state: *δ*_*i*_ for state 1^(*i*)^, *ω*_*i*_ for state 2^(*i*)^ without therapy and *α*_*i*_ for state 2^(*i*)^ with therapy.

3 A healthy individual remains in state 0 and thus collects a payoff of 1 indefinitely. Taking into account time-discounting, this person has a payoff of 1*/ρ* = 20.

4 Note that this does not mean that subsequent rounds of therapy offer no benefits as patients under therapy have a longer life expectancy then those who are not even if *λ*_*i*_ = 0.

5 Note that at state 2^(*i*)^ the patient makes a decision on the *i* + 1th round of treatment.

6 The expected time spent in state 1^(*i*)^ is 1*/δ* = 10 in this example, while toxicity level upon leaving state 1^(*i*)^ if it was at level *z*′upon entering it is *z*′ *δ/*(*δ* + *ζ*), so an initial toxicity level of around 0.5 decreases to around 0.38.

7 In Example 3.5’s parametrization, the bias in 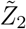 is around 0.005, amounting to 1% of 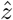 with the estimate being lower, hence the second waiting time is slightly underestimated; the first waiting time’s toxicity is unaffected by the bias, while all subsequent rounds have barely measurable payoff-effects.

